# *In vivo* efficacy of fidaxomicin against *rpoB* mutant *Clostridioides difficile* infection

**DOI:** 10.1101/2025.04.17.649461

**Authors:** Mai Thu Hoai, Yutaro Hitomi, Tsutomu Fujii, Yoshitomo Morinaga

## Abstract

**Objectives:** *Clostridioides difficile* infection (CDI) is a well-known healthcare-associated diarrheal disease. Fidaxomicin, a key antibiotic used to treat CDI, targets *rpoB*. However, some clinical isolates have mutations in *rpoB*, which reduces their susceptibility to this antibiotic. In this study, the effects of *rpoB* mutations on the virulence of *C. difficile* and efficacy of fidaxomicin against CDI were evaluated *in vivo*.

**Methods:** An *rpoB* mutant strain (*C. difficile* G1073R-2024) with reduced fidaxomicin susceptibility was generated through spontaneous induction in a murine CDI model from the parental strain *C. difficile* VPI 10463. The virulence and therapeutic responses of the mutant strain were compared with those of the parental strain using a CDI model, including survival rate, body weight changes, clinical scores, and bacterial loads in feces.

**Results:** *C. difficile* G1073R-2024 had an amino acid alteration in Gln1073Arg and the minimum inhibitory concentration of fidaxomicin was >64 μg/mL. *In vivo* virulence was not significantly different between strains. Fidaxomicin treatment resulted in 100% survival rates and a comparable reduction in the bacterial load for both strains.

**Conclusions:** Fidaxomicin was effective against CDI caused by the *rpoB* mutant strain. The emergence of such mutations highlights the need for ongoing surveillance of drug resistance trends in clinical settings.

## INTRODUCTION

*Clostridioides difficile* is a gram-positive, obligate anaerobic bacterium that spreads via the fecal-oral transmission of resilient endospores. It is the leading cause of hospital-associated diarrhea and a major nosocomial pathogen [1]. *C. difficile* produces toxins, including toxin B (tcdB), that damage the colonic epithelium [2]. Alterations in the gut microbiota due to antibiotic exposure are widely recognized as principal risk factors for *C. difficile* infection (CDI).

Fidaxomicin, an antibiotic used to treat CDI, inhibits the RNA polymerase of *C. difficile*, thereby preventing its transcription. It is a key drug in recurrent and severe cases of CDI; however, some clinical isolates carry *rpoB* mutations in the β-subunit or the β′-subunit of RNA polymerase [3] that reduce drug susceptibility [3-5]. Although most previous studies have focused on the *in vitro* emergence of *rpoB* mutations [6-8], one study has isolated mutants from clinical samples and assessed their effects on spore germination and toxin production [4]. Although *rpoB* mutations in RNA polymerase subunits have been associated with reduced fidaxomicin susceptibility in clinical isolates, the potential of such mutations to cause treatment failure has not been fully evaluated. Previous studies have primarily relied on *in vitro* susceptibility testing, and few have explored the in vivo consequences of these mutations in the context of *Clostridioides difficile* infection (CDI) [6-8]. In this study, the virulence of an *rpoB* mutant strain and the efficacy of fidaxomicin against *C. difficile* infection caused by this mutant strain were evaluated using an *in vivo* model. By characterizing the in vivo behavior of the mutant, we aimed to determine whether reduced fidaxomicin susceptibility due to the *rpoB* mutation compromises treatment efficacy, thereby providing deeper insights into the clinical relevance of resistance-associated mutations and informing future surveillance and treatment strategies. Our study does not aim to characterize multiple clinical strains or compare the effects of different *rpoB* mutations. Instead, we focus on a specific *rpoB* mutant strain, isolated following fidaxomicin exposure in vivo, which has not been described in clinical settings. By evaluating the phenotype of this single, well-characterized mutant strain in both treated and untreated mice, our goal is to determine whether this *rpoB* mutation alone is sufficient to impact CDI severity or alter the therapeutic outcome of fidaxomicin treatment. This targeted approach helps bridge the gap between observed resistance mutations and their potential clinical relevance.

## MATERIAL AND METHODS

### Organisms

*C. difficile* strain VPI 10463 [9-11] was obtained from the American Type Culture Collection (ATCC, Manassas, Virginia, USA) and *C. difficile* G1073R-2024, which carries a G1073R mutation in the *rpoB* gene, was generated from the *C. difficile* VPI 10463 strain through in vivo selection following fidaxomicin treatment.

### Animal

Specific-pathogen-free (SPF) C57BL/6J male mice, aged 6–8 weeks and weighing 18–22 g, were purchased from Charles River Laboratories, Japan, Inc. Mice were housed under standard laboratory conditions (22 ± 2□°C, 50–60% humidity, 12-hour light/dark cycle) with free access to food and water. All mice were acclimatized for at least 3 days before experiments. All animal procedures were approved by the Institutional Animal Care and Use Committee of Toyama University (approval no. A2020med-18). These mice were also used in all later in vivo experiments throughout the study, unless stated otherwise.

### Induction of a fidaxomicin-resistant strain

*C. difficile* with elevated minimum inhibitory concentration (MIC) values against fidaxomicin was spontaneously induced in the CDI model mice. After the inoculation of cefoperazone-treated mice with VPI 10463, they were treated with a single oral dose of 30 mg/kg fidaxomicin. To isolate *C. difficile* strains, fecal samples were plated on taurocholate cefoxitin-cyclomerized mannitol agar. The isolates were then transferred to brain heart infusion media (Becton, Dickinson and Company, Franklin Lakes, NJ, USA) supplemented with L-cysteine (1 mg/mL) and yeast extract (5 mg/mL) (BHIS) agar containing fidaxomicin at concentrations of 2, 4, 8, 16, 32, 64, and 128 µg/mL. Colonies that grew on these plates were selected and the mutation location was identified by *rpoB* sequencing.

### rpoB sequencing and analysis

Sequencing was performed on *C. difficile* isolates with elevated MICs of fidaxomicin. DNA was extracted from colonies using 100 µL of 0.05 M NaOH, followed by heating at 95°C for 20 min and neutralization with 11 µL of 1 M Tris-HCl. The *rpoB* gene was amplified using AmpliTaq Gold 360 (Thermo Fisher Scientific, Waltham, MA, USA) and primers (Primer F: 5′– GATGCTCTTGAAGAAGCT–3′; Primer R: 5′–CAACATCTAGCTCAAATTCACC–3′). PCR conditions were initial denaturation at 95°C for 10 min, followed by 40 cycles of 95°C for 30 s, 56°C for 30 s, and 72°C for 1 min, with a final extension at 72°C for 7 min and storage at 4°C. The PCR products were purified using EnzSAP (Edge BioSystems, San Jose, CA, USA), and the labeling reaction was performed using a SupreDye Cycle Sequencing Kit (Edge BioSystems). The labeled products were purified using a BigDye Xterminator Purification Kit (Thermo Fisher Scientific). Sequencing was performed using an ABI 3500 Genetic Analyzer (Thermo Fisher Scientific), and the results were analyzed by comparison with the *C. difficile* VPI 10463 complete genomes (GenBank Accession: NZ_CM000604.1).

### Susceptibility testing to fidaxomicin

The MICs of fidaxomicin for parental and mutant strains were determined by agar dilution according to Clinical and Laboratory Standards Institute standards [12]. The MIC of fidaxomicin was determined after incubation in an anaerobic chamber at 37°C for 24 h. All assays were performed in triplicate.

### CDI model

Mice were pretreated with cefoperazone (0.5 g/L) in drinking water for 10–14 days to induce intestinal dysbiosis, followed by a 2-day recovery period with sterile water. Then, mice were challenged with two strains of toxin B-producing *C. difficile* spores, as previously reported [13]. CDI model mice were orally administered 30 mg/kg fidaxomicin twice daily for 3 days [14]. Changes in survival rate, body weight, clinical score, and number of *C. difficile* in feces were measured for at least 5 days or until the mouse died. The clinical scores were determined as previously reported [15].

### Spore preparation

*C. difficile* VPI 10463 and *C. difficile* G1073R-2024 were initially cultured in 2 mL of Columbia broth (Becton, Dickinson and Company) for 24 h under anaerobic conditions. The culture medium was then transferred to a Clospore liquid medium and incubated anaerobically at 37°C for 5–7 days. After incubation, spores were harvested by washing three to five times with sterile water to remove vegetative cells. The purified spores were stored at 4°C until further use.

### DNA extraction from feces

Fecal samples from mice were collected and stored at -20°C until DNA extraction. Genomic DNA was extracted using an Omega Bio-TEK Stool DNA Kit (Omega Bio-Tek, Norcross, GA, USA) following the manufacturer’s protocol. The extracted DNA was quantified and assessed for purity using a spectrophotometer before further analyses.

### qPCR

Quantification of the expression of *tcdB* and *16S rRNA* was performed using a LightCycler 96 system (Roche, Switzerland) with LightCycler 96 SW 1.1. The reaction was carried out in a two-step cycling program: initial denaturation at 96°C for 60 s, followed by 40 cycles of denaturation at 95°C for 10 s and annealing/extension at 60°C for 50 s. Relative expression of *tcdB* (Primer pair contains F: CCAAARTGGAGTGTTACAAACAGGTG, R: GCATTTCTCCRTTYTCAGCAAAGTA) was analyzed by comparing the two experimental groups using the ΔΔCt method, with *16S rRNA* as a reference (Primer pair contains 27F: AGAGTTTGATCMTGGCTCAG, 1492R: TACGGYTACCTTGTTACGACTT).

### Data analysis

All statistical analyses were performed using GraphPad Prism 10.1 (GraphPad Software, Boston, MA, USA). The choice of statistical tests was determined based on the data distribution and experimental design. For normally distributed data, unpaired *t*-tests were used for pairwise comparisons. In cases where multiple groups were compared, one-way or two-way ANOVA was applied, depending on the design of the experiment (e.g., for one-way ANOVA, one independent variable was tested across multiple groups; for two-way ANOVA, two independent variables were tested simultaneously). The assumption of normality was assessed using Shapiro-Wilk test and/or visual inspection of the data (e.g., histogram or Q-Q plot). If the data were found to be non-normally distributed, non-parametric tests such as Mann-Whitney U test (for pairwise comparisons) or Kruskal-Wallis test (for multiple group comparisons) were used instead. Post-hoc analyses were conducted using Dunnett’s multiple comparison test (for comparisons against a control group) or Sidak’s multiple comparison test (for specific group-wise comparisons). All p-values were adjusted for multiple comparisons to control for type I errors. Data are presented as mean ± standard deviation (SD) or mean ± standard error of the mean (SEM), as indicated in figure legends. Statistical significance was set at p < 0.05, and a p-value < 0.05 was considered statistically significant.

## RESULTS

### Sequencing of C. difficile strains with reduced susceptibility to fidaxomicin

Only one isolate showing stable growth was obtained after the spontaneous induction of the fidaxomicin-resistant strain. This strain had an incidental mutation in the *rpoB* gene, characterized by a nucleotide substitution at position A3221G, resulting in a Q1073R amino acid replacement. This mutant strain, *C. difficile* G1073R-2024, exhibited MICs (64 μg/mL) significantly lower susceptibility to fidaxomicin than the parental strain (0.25 μg/mL).

### Phenotype of CDI caused by the rpoB mutant strain

The phenotypic characteristics of the mutant strain were compared with those of the parent strain in a CDI model (Fig. 1A). In this model, the parental strain *C. difficile* VPI 10463 caused lethal disease, and all mice died within 48 h of inoculation, while the mutant strain also exhibited a lethal phenotype, and the survival rate was like that of *C. difficile* VPI 10463 (Fig. 1B). There were no significant differences in body weight change (Fig. 1C) or clinical score (Fig. 1D) between groups. The bacterial load in the feces 48 h post-inoculation was also not significantly different between strains (Fig. 1E).

**Figure 1.**
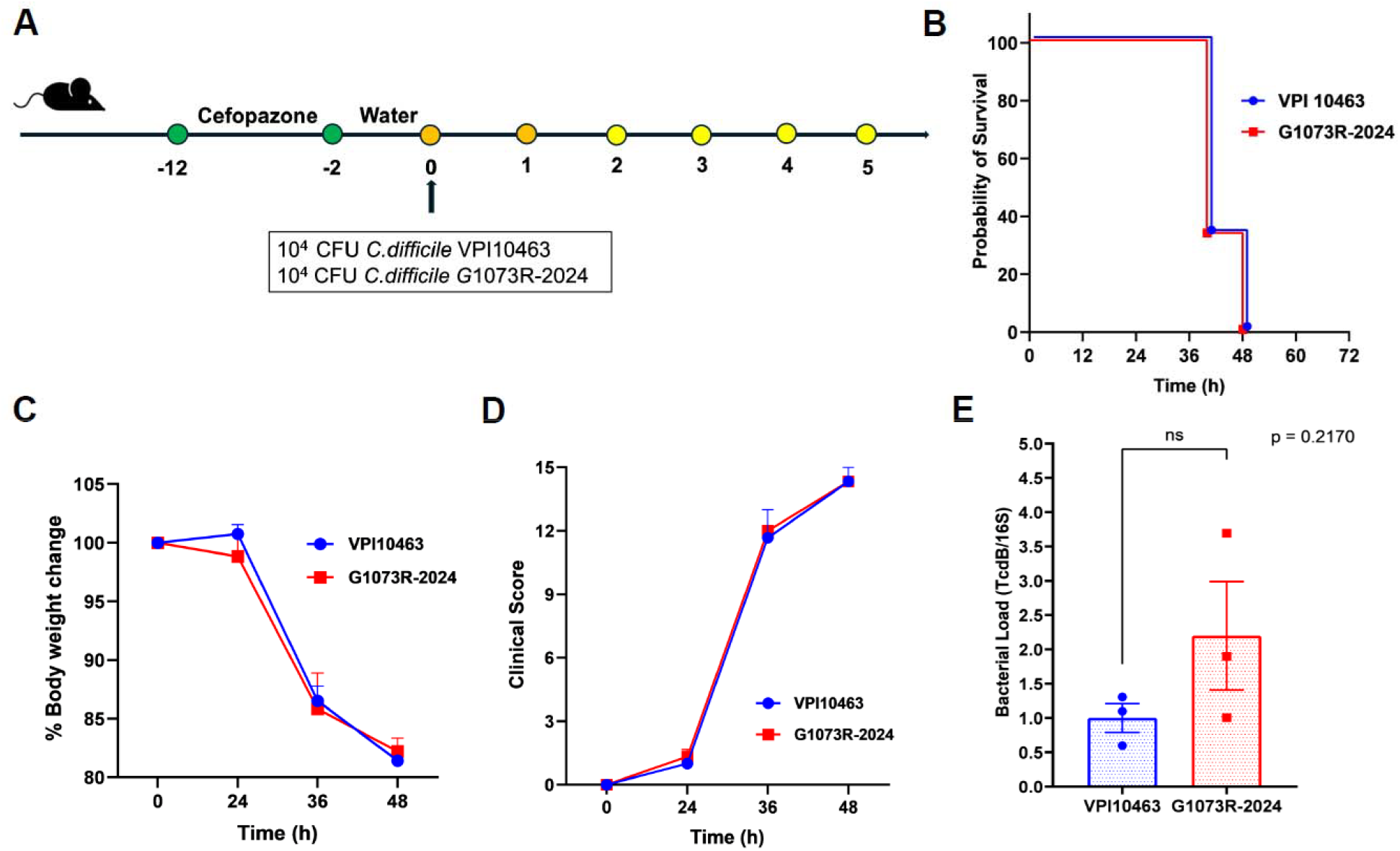
*rpoB* mutant strain caused the phenotype of CDI. **A**. Schematic of experimental procedure (n = 3, each group). **B**. Survival rate post-inoculation. **C**. Body weight change. **D**. Clinical score. **E**. Bacterial load in feces 2 days post-infection. Error bars represent the standard error of the mean.

### Efficacies of fidaxomicin treatment against the rpoB mutant strain

The therapeutic effects of fidaxomicin were compared between the parental and *rpoB* mutant strains in the CDI model (Fig. 2A). After 3 days of fidaxomicin treatment, both groups showed 100% survival on day 5 (Fig. 2B), with no significant differences in body weight changes (Fig. 2C) or clinical scores (Fig. 2D). The bacterial load in the feces on day 5 was also not significantly different between groups (Fig. 2E).

**Figure 2.**
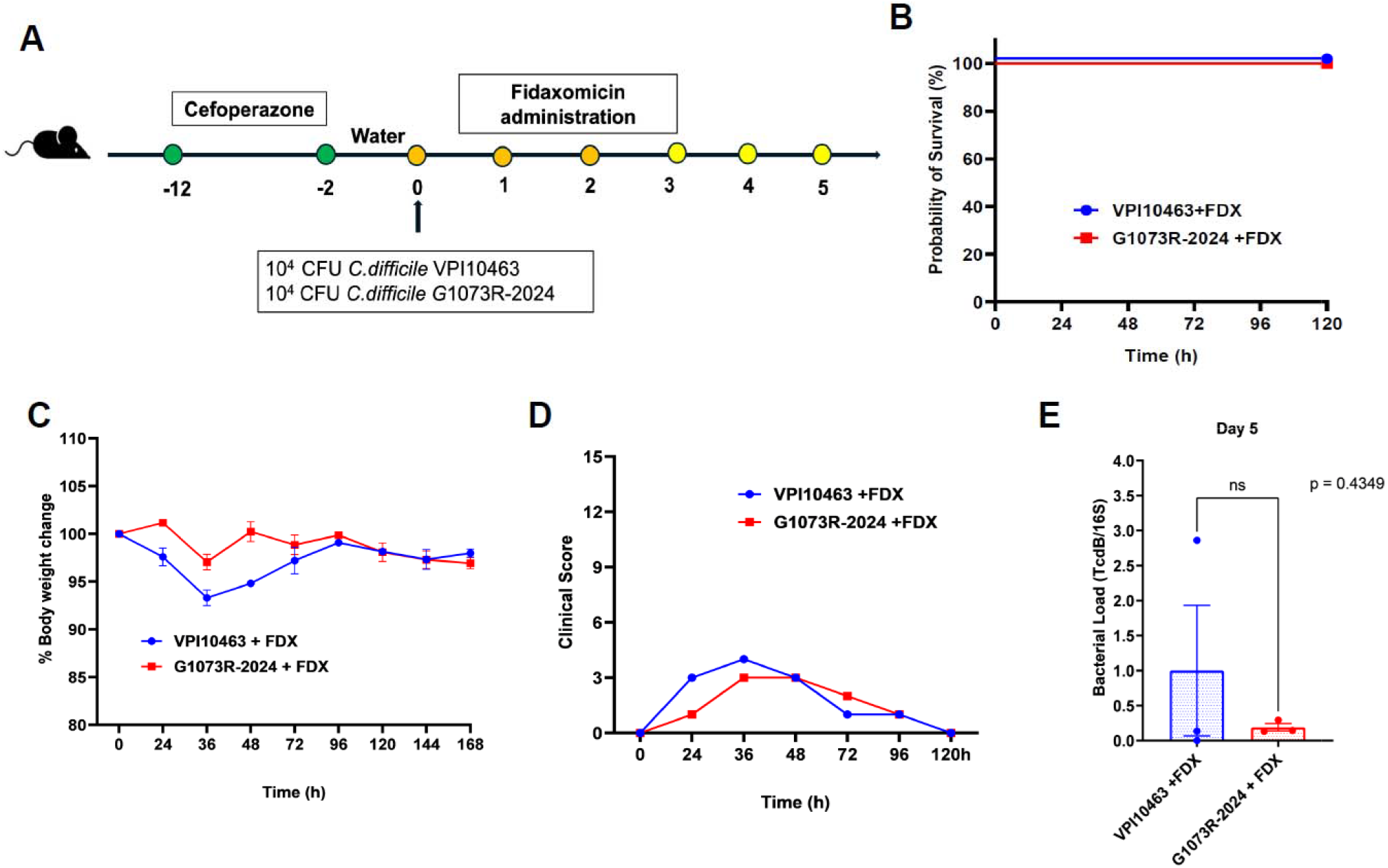
Efficacies of fidaxomicin treatment against *rpoB* mutant strain. **A**. Schematic of experimental procedure (n = 3, each group). **B**. Survival rate after fidaxomicin treatment. **C**. Body weight change. **D**. Clinical score. **E**. Bacterial load in feces 5 days post-infection. Error bars represent the standard error of the mean.

## DISCUSSION

In the present study, we examined the effect of *C. difficile rpoB* mutation on the virulence and efficacy of fidaxomicin *in vivo*. Fidaxomicin showed efficacy against CDI caused by *rpoB* mutant strains. We found no significant differences in virulence between the *rpoB* mutant strain and its parental counterpart *in vivo*. This finding contradicts a previous report that *rpoB* mutations affect bacterial fitness, including growth, sporulation, and toxin production, leading to reduced virulence *in vitro*. This discrepancy may be attributed to differences in the mutation sites within the *rpoB* gene; previous studies have focused on the V1143 mutant strain [8], whereas this study investigated the Q1073 mutant strain. Our findings suggest that *C. difficile* can acquire fidaxomicin resistance through *rpoB* mutations without compromising bacterial fitness. This conclusion is supported by our in vivo results showing that the rpoB mutant strain retained virulence comparable to the parental strain, as indicated by similar survival rates and fecal bacterial loads. Moreover, fidaxomicin remained effective against CDI caused by the mutant strain, indicating that while resistance was present, it did not render the treatment ineffective. Together, these data emphasize that certain resistance-associated *rpoB* mutations may not impair pathogenicity, underscoring the importance of continued surveillance and functional characterization of clinical isolates with reduced fidaxomicin susceptibility.

In addition to evaluating fidaxomicin efficacy, the virulence of the *rpoB* mutant strain was also assessed. Our vivo findings demonstrated that the mutant strain retained high virulence, as evidenced by the rapid onset of severe disease and mortality comparable to the parental strain VPI 10463. This observation suggests that the *rpoB* mutation conferring reduced susceptibility to fidaxomicin does not result in an attenuated phenotype.

Previous studies have shown that mutations in *rpoB* can affect bacterial fitness and virulence in various pathogens, such as Mycobacterium tuberculosis and Staphylococcus aureus; however, these effects are often context- and strain-dependent [16-19]. In *C. difficile*, evidence on how *rpoB* mutations influence virulence is still limited. Our results contribute to this gap by indicating that, at least in the case of the mutation studied here, the virulence remains unchanged. These findings highlight the potential clinical relevance of *rpoB* mutant strains, as they may both resist fidaxomicin treatment and retain full pathogenic capacity.

Fidaxomicin was effective against *rpoB* mutant strain-induced CDI model. One possible explanation for this is the pharmacokinetic profile of fidaxomicin. Fidaxomicin can achieve exceptionally high and prolonged fecal concentrations, with mean levels exceeding the MIC 5,000-fold [20, 21]. Although the strain used in the present study is a well-known and representative substitution related to susceptibility to fidaxomicin [3, 9], other substitutions in *rpoB* (e.g., D814Y [22], V1143F [8,12], V1143G [4, 12, 22, 23], V1143D [3, 8,12, 24], V1143N, and V1143L [4, 23]) have been also linked to reduced fidaxomicin susceptibility. However, the MIC values of fidaxomicin for these strains were comparable to or lower than those for our strain. Therefore, fidaxomicin may be effective against CDI caused by other *rpoB* mutant strains.

A noteworthy feature of our study is the spontaneous emergence of this Q1073R mutation *in vivo* because the same mutation had previously been identified only *in vitro* [5,12]. These findings suggest that *rpoB* mutants are prone to emerging *in vivo*. Therefore, analyzing trends in susceptibility to fidaxomicin and the underlying genetic mutations in isolates from patients after treatment with fidaxomicin should be continued.

This study had several limitations. First, our findings are based on a single *rpoB* mutant strain, which may not fully represent the diversity of clinically reported *rpoB* mutations [8]. Second, we did not assess the emergence of secondary mutations during or after treatment.

In conclusion, *in vivo*, virulence was not different between parental and mutant strains and fidaxomicin was effective against CDI caused by the *rpoB* mutant strain. Thus, our findings suggest the potential for the continued use of fidaxomicin against infections caused by *C. difficile* strains exhibiting resistance. However, this does not negate the need for ongoing surveillance regarding the emergence of drug-resistant mutant strains in clinical settings. Future studies should explore the impact of different *rpoB* mutation sites on bacterial fitness, virulence, and treatment outcomes, as well as assess the prevalence and clinical relevance of such mutations in patient populations.

## TRANSPARENCY DECLARATION

### Data availability

The authors confirm that the data supporting the findings of this study are available in the article.

### Declaration of Generative AI and AI-assisted technologies in the writing process

During the preparation of this study, the authors used Perplexity to improve readability and language. After using this tool, the authors reviewed and edited the content as needed and take full responsibility for the content of the publication

